# Is M1-L121E a good mimic on microbial rhodopsin? A viewpoint from excited-state dynamics

**DOI:** 10.1101/2023.11.03.565439

**Authors:** Gaoshang Li, Jiajia Meng, Shuang Yu, Xiaolu Bai, Jin Dai, Yin Song, Xubiao Peng, Qing Zhao

## Abstract

Microbial rhodopsin, an important photoreceptor protein, has been widely used in several fields, such as optogenetics, biotechnology, and biodevices *etc*. However, current microbial rhodopsins are all transmembrane proteins, which both complicates the investigation on the photoreaction mechanism and limits their further applications. Therefore, a suitable mimic for microbial rhodopsin can not only provide a better model for understanding the mechanism, but also can extend the applications. The human protein CRABPII turns out to be a good template for design mimics on rhodopsin, due to the convenience in synthesis and the stability after mutations. Recently, Geiger *et al.* designed a new CRABPII-based mimic M1-L121E on microbial rhodopsin with the correct 13-cis (13C) isomerization after irritation. However, it still remains a question how similar it is compared with the natural microbial rhodopsin, in particular in the aspect of the photoreaction dynamics. In this article, we investigated the excited-state dynamics of this mimic by measuring its transient absorption spectra. Our results reveal that there are two components in the solution of mimic M1-L121E at PH=8, known as protonated Schiff base (PSB) and unprotonated Schiff base (USB) states. In both states, the photoreaction process from 13-cis (13C) to all-trans (AT) is faster than that from the inverse direction. In addition, the photoreaction process in PSB state is faster than that in the USB state. In the end, we compared the isomerization time of the PSB state with the properties of the microbial rhodopsin, and confirmed that the mimic M1-L121E indeed captures the main feature of the rhodopsin and is a good model of microbial rhodopsin in the photoreaction dynamics. However, our results also reveal significant differences in the excited-state dynamics of the mimic relative to the natural microbial rhodopsin, including the slower PSB isomerization rates in both 13C-AT and AT-13C directions, as well as the unusual USB photoreaction dynamics at PH=8. Such unique properties have not been observed in the natural rhodopsin, which could further deepen the understanding in photoreaction mechanism of the photosensitive proteins.

## Introduction

Rhodopsin, an important photoreceptor, is widely distributed in microbes and animals,^1,2^ where it is categorized as type-I and type-II, respectively. Unlike animal rhodopsin, which mainly participates in signal transduction during visualization, microbial rhodopsin has diverse functions, including ion pumps, sensors, channels, enzymes, phototaxis, and even the regulation of gene expression. Microbial rhodopsin can be found in several microorganisms, including archaea, bacteria, and Eukarya. Based on the information from its origin species and sequence similarities, microbial rhodopsin can be further classified into bacte-riorhodopsin (BR),^3–9^ proteorhodopsin,^10–12^ Anabaena sensory rhodopsin (ASR),^13–15^ viral rhodopsin,^16^ schizorhodopsin (SzR)^17,18^ and Heliorhodopsins (HeR)^19–21^ *etc*.

Retinal proteins are characterized by fast photoisomerization that occurs within femtoseconds to picoseconds with a high quantum yield, ^22–25^ which highlights its efficiency as a molecular machine for converting solar energy into chemical energy. Microbial rhodopsin has been applied in several fields, such as optogenetics, biotechnology, and biodevices, etc.^26–33^ Despite the successful applications of microbial rhodopsin, several questions regarding the photoreaction mechanisms, such as how it regulates the absorption spectrum, the factors affecting the isomerization rate, and the photoinduced protonation/deprotonation processes remain unresolved. The large size of microbial rhodopsin and the various time scales of the entire photoreaction process make these questions challenging to answer. In particular, the presence of transmembrane regions in rhodopsin makes it even more difficult to study.

Hence, a rhodopsin mimic would be beneficial for understanding the experimental and theoretical aspects of the photoreaction mechanism. So far, several rhodopsin mimics have been designed based on CRABPII or CRBPII,^34–42^ and their dynamics have been well investigated.^43–46^ In these mimics, the chromophore retinal can covalently bond to the protein by forming a Schiff base (SB) with the Lys residue, which is essential for the photoreaction process. However, unlike the traditional natural rhodopsin that consists of seven transmembrane *α*-helices connected by extracellular and intracellular loops, these mimics are not membrane proteins but have *β*-barrel-like structures with only half the number of residues seen in natural rhodopsin. Another significant difference is that the retinal in these mimics can only adopt an all-trans conformation. Only recently, Geiger et al. designed a new variant M1-L121E based on the human protein CRABPII to effectively mimic microbial rhodopsin.^47^ They showed that the single crystal form of the mimic had a similar photocycle as the natural microbial rhodopsin. However, the excited state dynamics of this mimic have not been analyzed yet. Hence, the similarity in the aspect of excited-state dynamics between the mimic and the microbial rhodospin remains a question.

In this article, we used the transient absorption spectrum (TAS) with different pumping sources to investigate the excited state dynamics of the mimic M1-L121E in solution. We revealed that the isomerization of the 13-cis structure is faster than the all-trans structure when the SB is in a protonated state, which is similar to microbial rhodopsin.^48–52^ However, the isomerization rates of both directions are much slower in M1-L121E compared to natural microbial rhodopsin. This difference leads to the weak fluorescence in the rhodopsin mimic, which is hardly observed in natural rhodopsin. Interestingly, in M1-L121E, the SB is always in a mixture of protonated and unprotonated states that co-exist at pH = 8. The absorption spectra of both states heavily overlap, which has not been observed in natural microbial rhodopsin or the previous mimics. As a result, the absorption spectrum of M1-L121E shows different behaviors under light irritation. In summary, by studying the excited-state dynamics of the microbial rhodopsin mimic M1-L121E, we show that the engineered mimic possesses the main features of natural microbial rhodopsin in the subnanosecond time scale dynamics. However, the mimic also has unique properties, which provide valuable hints for future microbial rhodopsin engineering, such as to design proteins that can response to the light irritation in a wide wavelength band.

## Results and discussion

### Protein Structures and the Steady Absorption Spectrum

Figure 1A displays the X-ray crystallographic structure of the protein and retinal chromophore in M1-L121E separately, retrieved from the Protein Data Bank (PDB). When the retinal binds to the protein, it can form either all-trans (AT) or 13-cis (13C) state as shown in Figures 1B and 1C, respectively. According to the previous X-ray crystallographic observation,^47^ the two structures are generated from dark adaption and UV irritation, which we refer to as dark and UV states, respectively throughout this paper.

**Figure 1:**
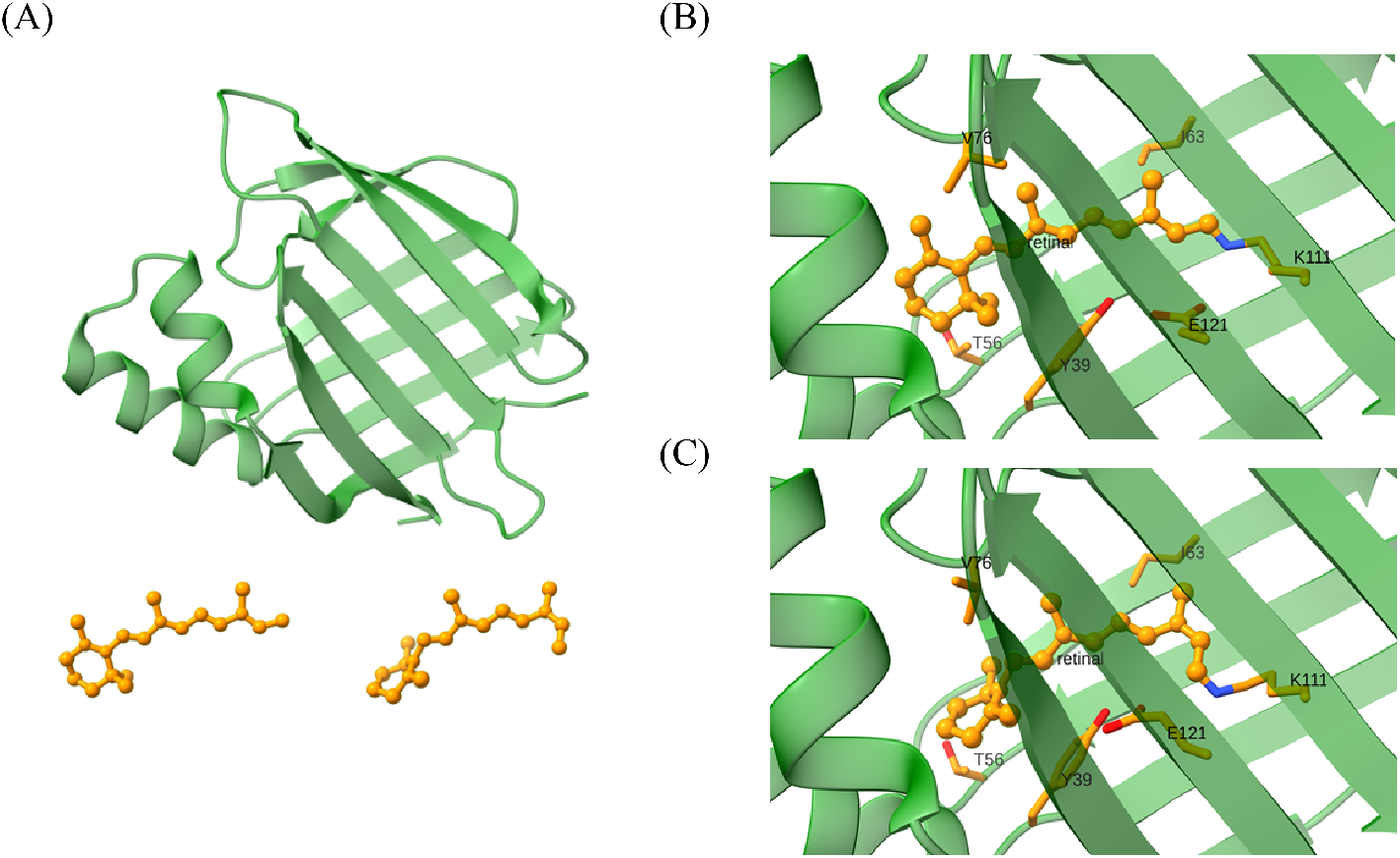
The structure of the microbial rhodopsin mimic M1-121E.The protein is shown in the cartoon representation. (A) The structures of the components protein and retinal in the mimic. (B) The close-up view in the chromophore binding region around the all-trans retinal chromophore. (PDB 6MOP)^47^ (C) The close-up view in the chromophore binding region around the 13-cis retinal chromophore. (PDB 6MQZ)^47^

In Figure 2A, the measured steady absorption spectra of M1-L121E have been shown as blue and red curves for the dark and UV states, respectively. Each curve has two peaks, corresponding to the protonated and unprotonated Schiff bases (PSB and USB). Hence, we conclude that in M1-L121E, the PSB and USB states co-exist at pH = 8, irrespective of the dark or UV states. Through decomposition of the absorption spectral using Gaussian functions, we further identified the peaks of the maximum absorption for different protonation states in both dark and UV states (Fig2 B and 2 C, respectively). As shown in Figure 2 B, the USB and PSB are located at 374 nm and 470 nm, respectively in the dark state, but in the UV state, they are slightly blue-shifted to 361 nm and 468 nm. Further on, we noticed that the decomposed spectra components of USB and PSB overlap significantly in the range of 370 nm-450 nm for the dark state and 370 nm-440 nm for UV state, indicating that the light in this band would activate both USB and PSB.

**Figure 2:**
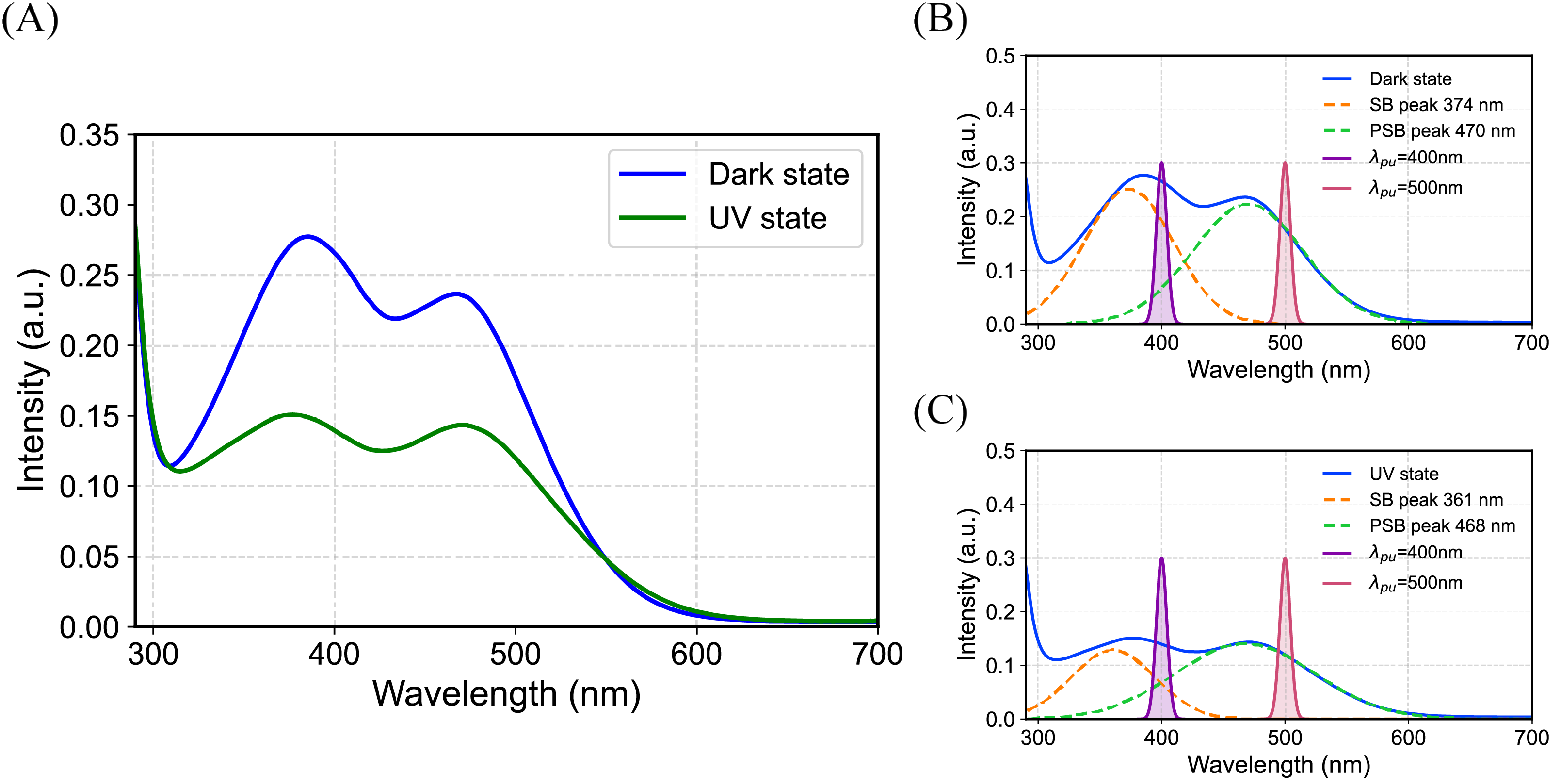
(A) The steady absorption spectra for the Dark (blue solid line) and UV states (green solid line) of M1-121E. (B) The decomposition of the steady absorption spectra for the Dark state and (C) UV state. The absorption spectra for the SB (orange dashed line) and the PSB (green dashed line) undergo double Gaussian fitting.

Notably, the absorption of both SB and PSB in the UV state is significantly decreased compared to the dark state in our experiment, consistent with a previous study.^47^ This study suggested that this might be because 13C-retinal has lower extinction coefficients compared to the AT state, causing a drop in the absorption after isomerization from the AT to the 13C state due to UV irritation. In our experiment, we further validated by measuring the absorption spectra at different dark adaption times after UV irritation. We observed that the absorption slowly increases toward the level of the dark state with time due to the thermal isomerization from 13C to AT states.

### The transient absorption spectral in UV and dark states

As shown in Figure 2, the system contains both PSB and USB components at pH 8 irrespective of the dark or UV states. According to the crystallographical result, the conformation of SB is mainly AT in dark state while becomes 13C in UV state. As a result, there are in total four states of SB in our system, known as AT-PSB, AT-USB, 13C-PSB and 13C-USB. In our experiment, 13C (UV) and AT (dark) states were controlled by UV irritation along TAS measurement, while the USB and PSB states are distinguished via different pumping lasers. As shown in Fig.2 B-C, only the PSB component was excited when the system was pumped with the 500 nm femtosecond laser. Hence, we use this laser to study the excited-state dynamics of PSB. However, for USB we could not obtain the pure TAS using the visible light pumping as its absorption spectrum in visible light region heavily overlaps with PSB’s. As an indirect way, we first measured the TAS of the mixed state with a pumping laser at 400 nm. Afterwards, we performed global analysis on the TAS of the mixed state using the constraints of the PSB dynamics that had been resolved with the 500 nm pumping laser. Finally, we obtained the excited-state dynamics of corresponding USB state.

#### TAS with 500 nm pump

Figure 3A and 3B show the TAS heatmaps for M1-L121E in the Dark and UV states, respectively. The corresponding absorption spectra at several different delayed time points are listed right each heatmap. We observed that the TAS for both states mainly peaked between 520 nm and 740 nm. The information is missing around 500 nm due to the scattering of the light from the pump. Moreover, although signals were present between 420–450 nm, the signal-to-noise ratio was too low to distinguish it from noise. Hence, we excluded these regions from our analysis.

**Figure 3:**
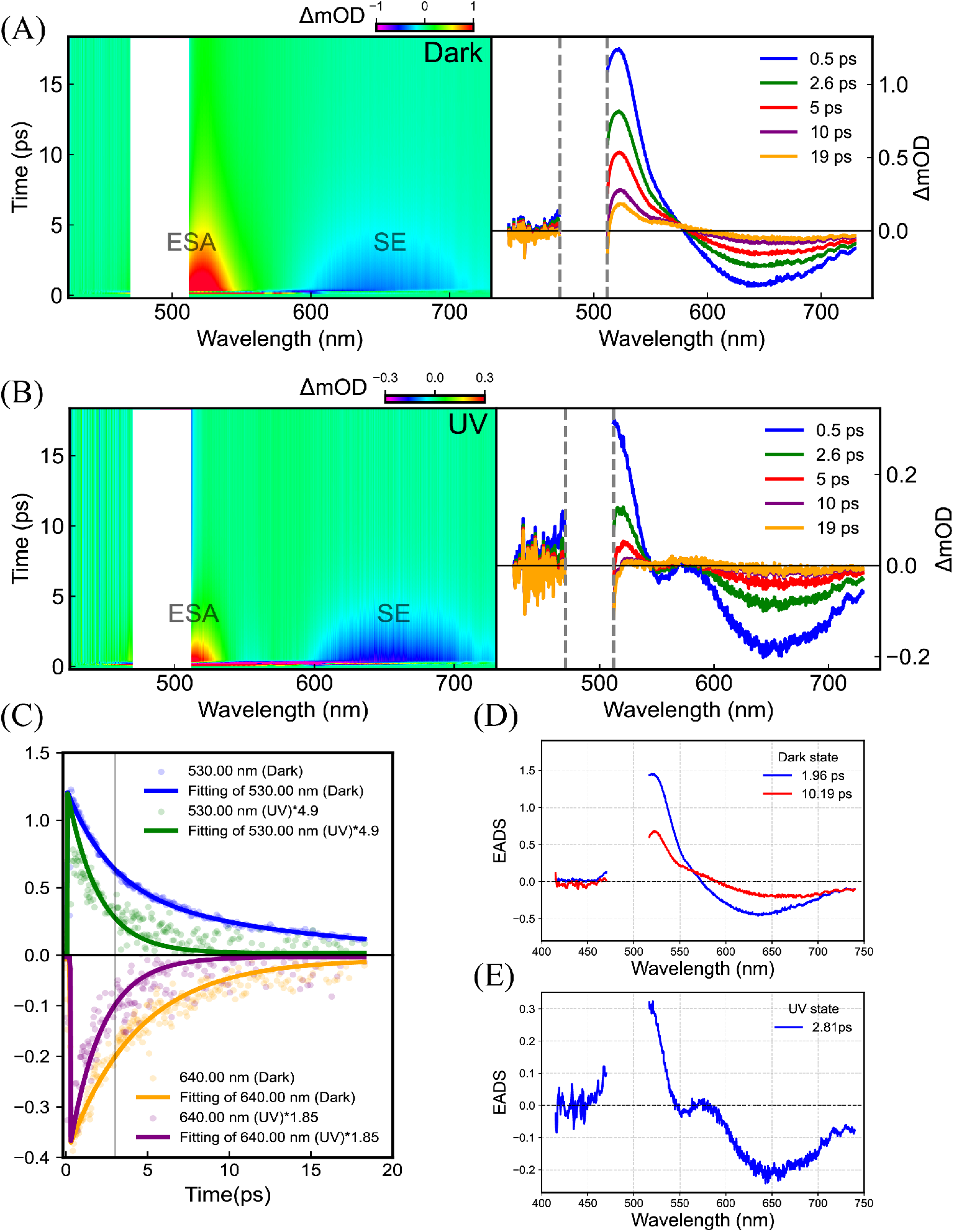
(A) The left displays TAS heatmap in the Dark state and (B) UV state. The right presents TAS curves at different time points. (C) Time traces of the Dark and UV states at 530 and 640 nm (circles) with traces reconstructed from the global fit (solid lines). The single wavelength data of 530 and 640 nm in the UV state are multiplied by 4.5 and 1.8 respectively for better comparison. (D) The EADS derived by global fitting on the TAS data for the Dark state and (E) UV state.

In the dark state, the TAS displays clear positive peaks in the 515–550 nm wavelength range, due to the excited-state absorption (ESA). We noticed a trend of very weak positive signals in the range of 450–470 nm, which might be due to the excited-state absorption of some residual USB components. The ground-state bleach (GSB) is invisible in our experiment due to the pump light scattering. Additionally, a broad region of stimulated emission (SE) can be observed within 600–700 nm. In the UV state, similar spectral features are observed, including the ESA around 450–470nm (weak signal) and 515–550 nm, and SE within the 600–700 nm range. However, by carefully comparing Figure 3A with 3B, we observed that the EAS peak in the UV state has a slight blue shift (4 nm) compared to that in the dark state. To facilitate a more comprehensive comparison, please refer to SIFig 2. Furthermore, the UV state also exhibits a stronger SE compared with the dark state.

To better understand the differences in the photoisomerization kinetics between the two states, we conducted exponential fitting analyses on the kinetics at several specific wavelengths. In Figure 3C, we applied an amplification factor to the UV state data for better visualization and observed that the decay rate in the UV state is notably faster. The results of single-wavelength fitting also demonstrate that the UV state can be accurately modeled by a single exponential decay, while the dark state requires a double exponential decay for a good fitting.

We also performed global analysis^53,54^ on the TAS data using a sequential model, where the initial excited state progressively transitioned into other states during its lifetime. For the dark state, three time constants were obtained for a good fitting (t1 *<* 0.15 ps, t2 = 1.96 ps, t3 = 10.19 ps). Notably, the shortest decay time, constant t1, was situated beyond the time resolution range of our measurements. This rapid decay can be influenced by coherence effects arising from factors such as the temporal overlap of pump and probe pulses, wave packet motion, and dynamic Stokes shift. Hence, we disregarded this ultrashort process and showed the EADS of the remaining two components in Figure 3D. For the EADS of the first component, the peaks of ESA and SE center at 530 nm and 630nm, respectively. After 1.96 ps, the reaction is transitioned to the second component, whose EADS shows a significant red-shifted in SE with a peak at 670 nm, indicating that the system has moved to another local minimum on the surface of the excited state. In the case of the UV state, we just obtained two time constants (t1*<* 0.15 ps, t2 = 2.81 ps) with which the TAS data had been fitted accurately enough. Again, as the first time constant is beyond the resolution of the equipment, it was disregarded. Figure 3E shows the EADS of the remaining component, which indicates the peaks of ESA and SE located at 530 nm and 650 nm, respectively.

Comparing the results of the global analysis on the TAS data with a 500 nm pump in the dark and UV states, we conclude that the UV state has one less process than the dark state, leading to its substantially faster decay. Interpreting from the viewpoint of the excited-state potential energy surface, the dark state will have an extra energy barrier than the UV state. Moreover, the SE process also differs depending on the states: In the dark state, there is a red shift in SE from 630 nm to 670 nm as the chemical reaction goes on, while it is always located around 650 nm during the whole photoreaction process of UV state.

#### TAS with 400 nm pump

We further investigated the dynamics of M1-L121E using the 400 nm pump light. Initial observations indicated that the dynamics are considerably slower, prompting us to employ a delay time of 100 ps for a more comprehensive investigation. Figure 4A and 4B show the TAS heatmaps of M1-L121E with the 400 nm pump in the dark and UV states, respectively. The corresponding absorption spectra at several different delayed time points are listed below each heatmap. Comparing Figure 4 with Figure 3, we observed that the time scale of the heatmap with the 400 nm pump is one order larger than that with the 500 nm pump for both dark and UV states, indicating that the photoreaction kinetics is significantly slower. As mentioned before, the system excited by 400 nm laser is in a mixed state with both PSB and USB, while the system excited by 500 nm is mainly in PSB state, we conclude that the photoreaction kinetics of USB is much slower than PSB.

**Figure 4:**
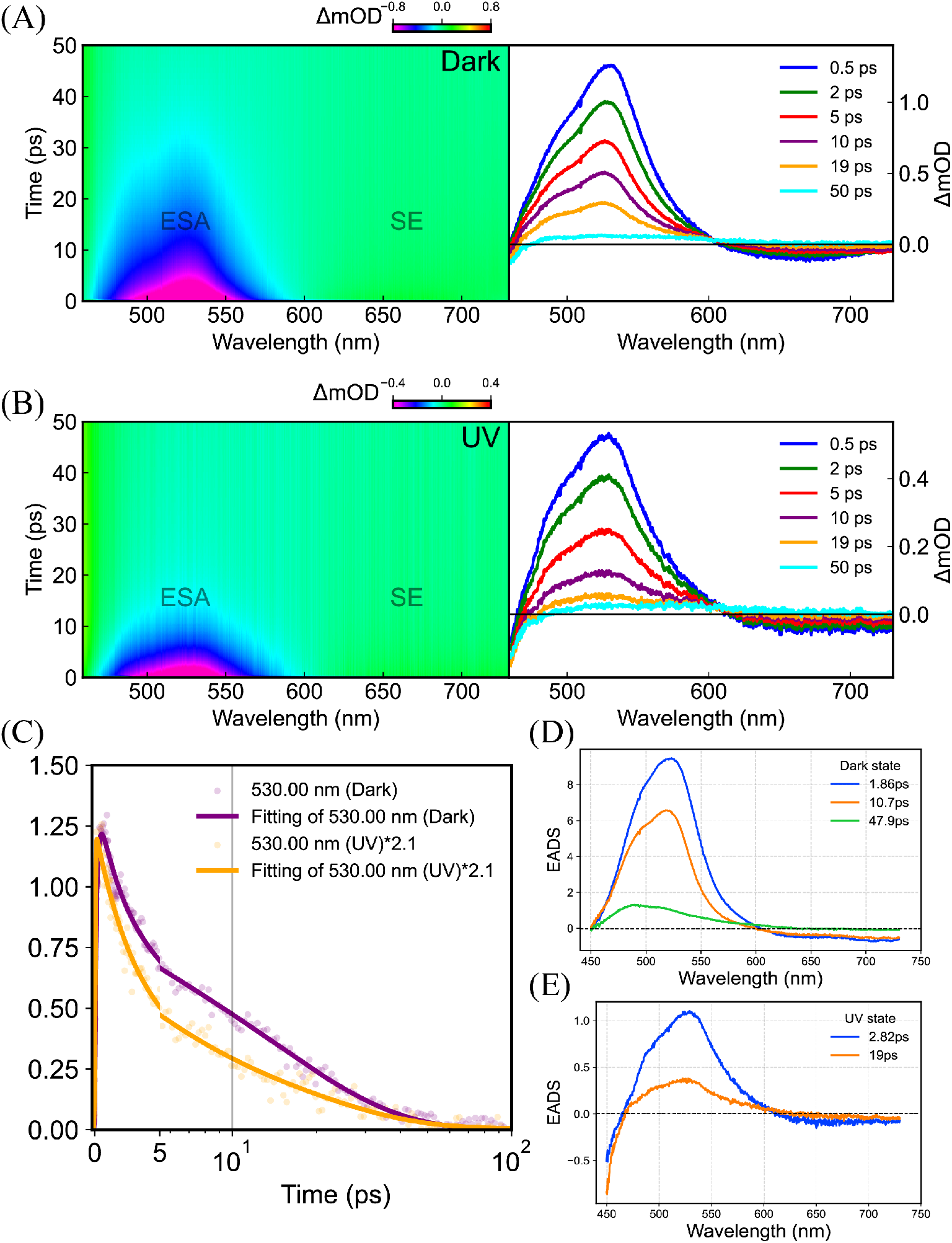
(A) The left displays TAS heatmap in the Dark state and (B) UV state. The right presents TAS curves at different time points. (C) Time traces of the Dark and UV states at 530 nm (circles) with traces reconstructed from the global fit (solid lines). The single wavelength data of 530 nm in the UV state are multiplied by 2.1 for better comparison. (D) The EADS derived by global fitting on the TAS data for the Dark state and (E) UV state.

From the heatmaps in Figure 4A and 4B, we observed that the TAS reveals positive and negative signals in the 450–600 nm and 600–700 nm ranges, respectively, for both states. These regions are attributed to ESA and SE, respectively. Notably, the EAS signals exhibit a noticeable blue shift with time going on in both states. We also observed a negative signal in the region shorter than 450 nm, indicative of the GSB of the sample. Although their TAS heatmaps share similar patterns, the kinetics in the UV state are much faster than in the dark state. Figure 4C show the fitting of single wavelengths at 530 nm for both states, providing a clear picture that the signal in the UV state decays faster than in the dark state. We performed the global analysis with the sequential model on the TAS heatmaps (Fig. 4). Since the system is in a mixed state with both PSB and USB, we added additional constraints during the analysis, including the time constants of the PSB kinetics, which have been resolved in the subsection “TAS with 500 nm pump.” Finally, we obtained three valid time constants (t1 = 1.86 ps, t2 = 10.7 ps, t3 = 47.9 ps) for the dark state, where the first two time constants were consistent to those of PSB, while the third one was unique for USB. Similarly, we obtained two valid time constants (t1 = 2.82 ps, t2 = 19 ps) for the UV state, where the first one is the same as that of PSB and the second is specific for USB. We note that there is an additional small time constant (*<*150fs) in above global analysis for both dark and UV states, but we have disregarded it taking the time resolution of our equipment into account.

The corresponding EADS profiles for the dark and UV states are shown in Figure 4D and 4E, respectively. For the dark state, the EADS profiles of the first two components contain the main features of the corresponding EADS profiles of PSB. For example, the EADS of the first component (t1 = 1.86 ps) has ESA and SE peaks at around 530 nm and 650 nm, respectively, similar to the corresponding EADS of PSB. However, it has an additional minor shoulder that was seen at 500nm in the ESA process, possibly due to the USB component. Similar behaviors were observed for the second component with a time constant of 10.7 ps. For the unique time constant t3 = 47.9 ps, its corresponding EADS has only positive signals but no negative signals, indicating this could be the production absorption process. The peak of the EADS profile was shifted to around 480 nm, corresponding to the absorption peak of the intermediate state. Similarly, the EADS profile of the first component (t1 = 2.82 ps) for the UV state (Fig. 4E) also captured the main features of the PSB, including ESA and SE peaks at 530 nm and 650 nm, respectively. Moreover, the EADS profile of the first component also showed an additional shoulder at 500 nm belonging to the ESA of the USB. The second EADS profile with a longer time constant belongs to USB only, where the ESA peaked at 525 nm and SE located around 650–700 nm.

In summary, we concluded that the time constant of the photoreaction kinetics of the USB was similar to that of PSB, barring an extra process with a longer lifetime in the picoseconds time scale. For the dark state, this extra process is related to the intermediate state on the potential energy surface of the ground state, while, for the UV state, it is related to the intermediate state on the potential energy surface of the excited state.

### Primary Photoreaction Dynamics of rhodopsin mimic M1-L121E

Based on the results of the global analysis on the TAS for USB and PSB in the dark and UV states, we could deduce the primary photoreaction dynamics of the microbial rhodopsin mimic M1-L121E at pH = 8 (Fig. 5).

**Figure 5:**
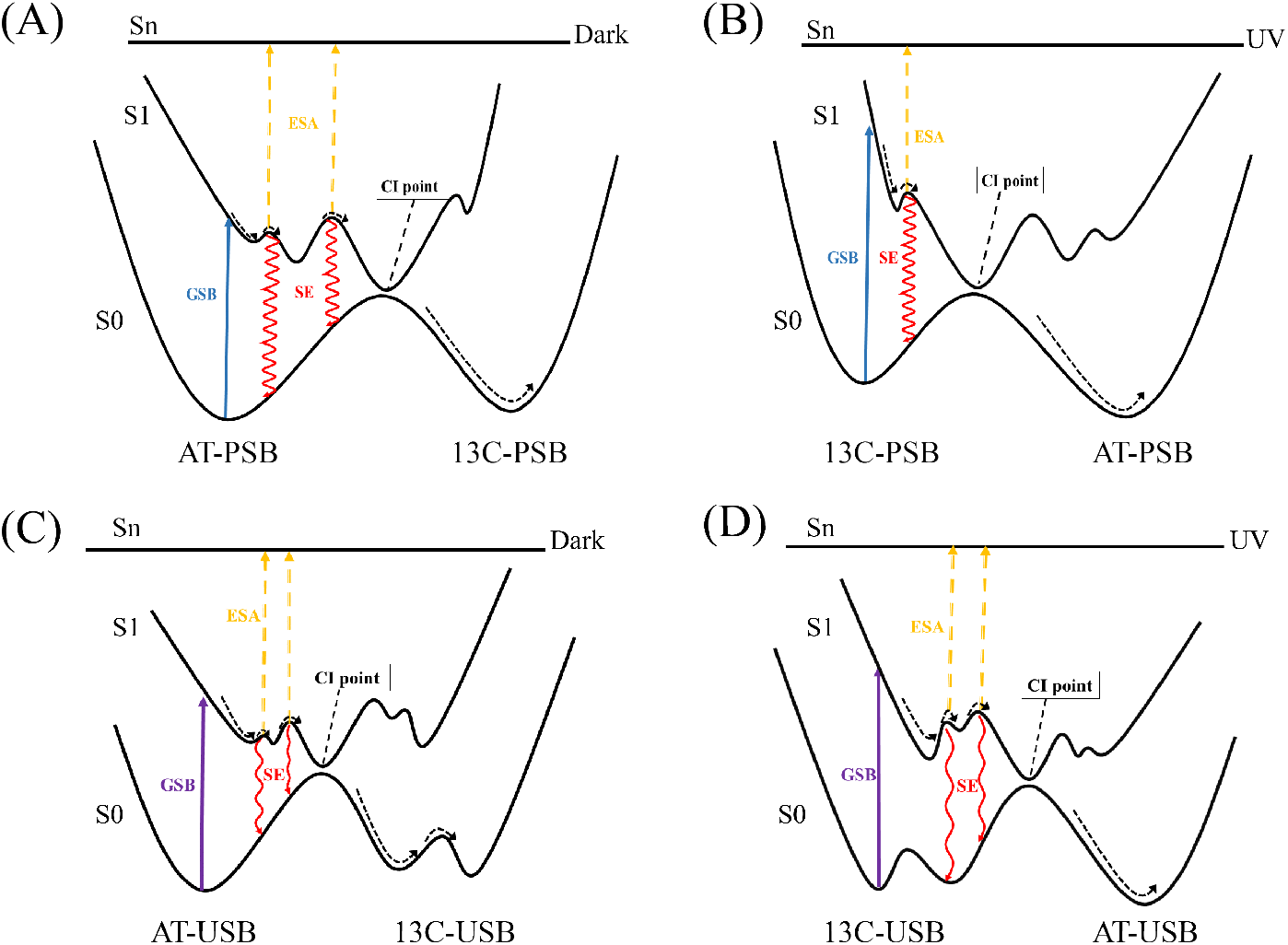
(A) Proposed reaction model outlining the primary events in the transition from M1-L121E in the all-trans (AT) state to the 13-cis (13C) state within the primary Schiff base (PSB) binding site. (B) The reverse transition from 13C-PSB back to AT-PSB state. (C) Transition from AT state to the unprotonated Schiff base (USB) state. (D) Reverse transition from 13C-USB back to AT-USB state.

In both the UV and dark states, the SB may undergo rapid structural or vibrational relaxation from the Frank-Condon state to its local energy minimum of the excited-state energy surface within 150 fs after photon absorption. Following this fast relaxation, distinct sequential chemical reactions for the dark and UV states start. In the case of PSB, when the system is in the dark state, the PSB first transits from the excited state of 13T–15T to an intermediate state on the excited-state energy surface (maybe 13C–15T or 13T–15C state) within 1.86 ps, then reaches the ground state of 13C-15C through the conical intersection point in 10.7 ps by overcoming an even higher energy barrier. If the system is in UV state, after overcoming a small energy barrier, the progression directly advances from the excited state of 13C-15C-PSB to the ground state of 13T-15T-PSB through the conical intersection point in 2.82 ps. In the case of USB, when the system is in the dark state, it needs to overcome similar energy barriers on the excited-state energy surface as the PSB in the dark state, passing through the conical intersection point to an intermediate state on the energy surface of the ground state, and finally reaching the more stable 13C15C ground state by overcoming an energy barrier, which takes 47.9 ps. If the system is in the UV state, it just needs to overcome two energy barriers on the excited-state energy surface to reach the ground state of 13T-15T USB, which takes 2.82 ps and 19 ps, respectively.

We note that the isomerization of retinal did not lead to significant changes in the protonation states in mimic M1-L121E, which can be seen from the comparison in the steady absorption spectral between dark state and UV state. If there is isomerization induced protonation, the absorption spectral for USB and PSB will show different behaviros after UV irritation. However, in our experiment, the absorption intensities of both USB and PSB in UV state get decreased compared with the dark state. Hence, we infer that the protation state is independent on the isomerization. According to Figure 5, we can see that the energy surface of the USB state is more complicated than the PSB state, leading to the longer photoreaction time with 400nm pumping. Although the photoreaction time in UV state is always shorter than in dark state for both USB and PSB, their detailed mechanisms are different. For PSB the reason is that the excited-state energy surface of the AT isomer is less smoother than 13C isomer, while for USB it is due to the existence of the intermediate state on the energy surface of the ground state before reaching the stable 13C isomer.

### Comparison with the microbial rhodospin

In this section, we compare the excited-state dynamics between the mimic and different kinds of microbial rhodopsin. In Table 1, we list the isomerization time constants of several different kinds of microbial rhodospin. We first note that the isomerization processes between 13C and AT happen in the protonated state (PSB) in all microbial rhodopsin. We observed that there are clear disparities in the dynamics between the transition from all-trans (AT) to 13-cis (13C) and the reverse transition from 13C to AT in some of microbial rhodopsins. For example, in BR, AntR and ASR, the isomerization processes from 13C to AT are much faster than those from AT to 13C.^49–52^ In our study, the exicted-state dynamics of the mimic M1-L121E reveal a similar property when it is in the pronated state (PSB). Through global analysis, we finally obtain the potential energy surface as a function of the isomerization reaction coordinates shown in Figure 5. We can see that the difference of the disparity is due to the presence of additional potential energy barrier on the 13C-PSB side of the the conical intersection point on the excited-state potential surface of the PSB, which is again similar to that of the microbial rhodopsins. However, we have to mention that the rate of the isomerization of the mimic is much slower than all the microbial rhodopsins in the table in either direction. This is not surprising since the microbial rhodopsins have been well optimized by the nature evolution. Unless the detailed mechanism of the photoisomerization of the rhodopsin is clear enough, it will be a great challenge to surpass the natural rhodospins in the aspect of the isomerization rate.

**Table 1:** Photoisomerization Time Constants of Diverse Microbial Rhodopsin.

| Microbial Rhodopsin | $\tau_{at-13c}$ | $\tau_{13c-at}$ |
| --- | --- | --- |
| BR<sup>48,55</sup> | $\sim 500$ fs | $\sim 200$ fs |
| PR<sup>56</sup> | $\sim 150$ fs | — |
| KR2<sup>57</sup> | $\sim 180$ fs | — |
| AntR<sup>51</sup> | $\sim 1.0$ ps | $\sim 300$ fs |
| ASR<sup>58,59</sup> | $\sim 750$ fs | $\sim 170$ fs |
| CaChR1<sup>60,61</sup> | $\sim 110$ fs | — |
| TaHeR<sup>62</sup> | $\sim 220$ fs | — |
| HeR 48C12<sup>62</sup> | $\sim 420$ fs | — |
| PoXeR<sup>63</sup> | $\sim 1.20$ ps | — |
| M1-L121E(PSB) | $\sim 12.2$ ps | $\sim 2.8$ ps |

Another great difference between the mimic and microbial rhodopsin lies in the fact that the mimic can automatically have the USB component at PH=8. Similarly, the USB also show that the photoreaction process in UV state is faster than that in the dark state. However, such a disparity is due to the additional energy barrier on the potential energy surface of the *ground state*, which is not observed in the microbial rhodopsin systems.

## Conclusion

Microbial rhodopsin is an important photosensitive protein. Designing a mimic for this protein provides a model to better understand the photoreaction mechanism in microbial rhodopsin, as well as extending the applications of microbial rhodopsin. Recently, Geiger et al. designed a microbial rhodopsin mimic M1-L121E and validate that the mimic has similar photocycles with microbial rhodopsin using the X-ray crystallographical measurements.^47^ However, as the excited-state dynamics of M1-L121E have not been investigated so far, it is yet to be ascertained if this mimic M1-L121E is a good model for microbial rhodopsin. In this study, we employed femtosecond transient absorption spectroscopy (FS-TA) to elucidate the photoreaction process of M1-L121E in the dark and UV states at pH = 8. Our findings demonstrated that M1-L121E has several remarkable properties similar to microbial rhodopsin in the excited-state dynamics when SB is protonated.In detail, we observed the dynamics of the transition from all-trans (AT) to 13-cis (13C) was different from that of the reverse transition from 13C to AT in microbial rhodopsin, as the latter was much faster. We also found some properties that were unique to M1-L121E. For example, M1-L121E possessed both PSB and SB states co-existing at pH = 8, while microbial rhodopsin is mainly in the PSB state. Further, we observed differences in the USB dynamics, mainly due to the energy barrier in the energy surface of the ground state. These phenomena are not observed in microbial rhodopsin.

In summary, our study shows that the properties of the excited-state dynamics of the rhodopsin mimic M1-L121E are similar to that of natural microbial rhodopsins, which further confirms that this mimic is a good model of microbial rhodopsin. Conversely, we also showed that M1-L121E possessed unique dynamics, which could be applied in some special situations.

## Experimental

### Sample preparation

All-trans-retinal, acquired from Sigma-Aldrich, was used as received without any additional purifications. As indicated previously,^42^ the protein-to-retinal equivalence ratio was maintained at 2:1 using a 1mM stock solution of retinal in ethanol. To prepare the dark state samples, we incubated the samples in the dark at room temperature for at least 12 hours. For measuring the transient absorption in the UV state, we used a 380 nm background light to continuously illuminate the sample to prevent thermal relaxation back to the all-trans structure. This helps keep the sample in a photo-quiescent state and ensures a significant accumulation of 13-cis structures. In the text, we refer to the former and latter as the ’Dark’ and ’UV’ states, respectively.

### UV-Visible Absorption Spectroscopy

Absorption spectra and slow kinetics were recorded at 25°C using a HORIBA UV-800C spectrophotometer at a spectral resolution of 1 nm. Dark-state samples were measured after a dark adaption period of 24 h. UV-state sample spectra were detected after a 5-min illumination with a UV-light-emitting diode (LED) (380 nm).

### Protein expression and purification

Samples were essentially prepared as described previously:^46^ M1-121E full-length sequences were cloned in a pET-28a plasmid with an N-terminal His-tag and expressed in Escherichia coli cells. We used codon-optimized E. coli constructs to maximize the efficiency and yield for protein purification. The target gene was transformed into Escherichia coli BL21 (DE3) cells (300 ng of DNA per 100 *µ*L of cell solution) following standard protocols and the cells were grown on Luria-Bertani (LB)-agar plates supplemented with antibiotics (Kanamycin: 50*µ*g/mL) at 37 *^◦^*C for 14 h. Then, 100 mL of LB medium containing 50 *µ*g/mL kanamycin was inoculated with a single colony and grown at 37 *^◦^*C with shaking overnight. The resulting culture was used to inoculate 1 L of LB containing 50 *µ*g/mL Kanamycin and was grown at 37 *^◦^*C while shaking till OD600 reached 0.8. The expression was induced with addition of isopropyl-*β*-D-thiogalactopyranoside (IPTG, Gold Biotechnology, 0.5mM) and the culture was shaken at 37 *^◦^*C for 4-5 h. The cells were harvested by centrifugation (4000 rpm, 15 min, 4 *^◦^*C) and resuspended in Tris-Hcl buffer (25 mM Tris, 150mM NaCl,pH=8.0,40 mL),disrupted by using a Ultra high pressure homogenizer.The solubilized membranes were isolated by high speed centrifugation (14000 rpm, 50 min, 4 *^◦^*C), and the supernatant was applied to a Ni-affinity column (HisTrap, GE Healthcare) at 4 *^◦^*C. After washing with Tris buffer, the mutants were eluted using a linear imidazole gradient. The purified protein was concentrated by centrifugation using an Amicon Ultra filter. The samples were further purified using Akta pure L with a HiTrap Desalting anion column (GE Health Sciences).

### Femtosecond Time-resolved Absorption Measurement

The experimental setup used was as described previously.^64^ It employed an 800 nm, 140 fs laser pulse generated by a Ti:Sapphire Regen amplifier as the primary light source to illuminate both TOPAS-C and a sapphire crystal. This laser pulse was divided into two beams, serving as the pump and probe pulses. To facilitate broadband white light probing, a portion of the laser output was focused onto a CaF_2_ plate.The pump pulses were directed through an optical parametric amplifier (TOPAS-C) to generate 400 nm and 500 nm excitation light at energies of 150 nJ and 200 nJ, respectively. To ensure precise control, we adjusted the polarization of the pump pulse to the ’magic angle’ (54.7°) relative to the horizontally polarized probe pulse.Both the pump and probe pulses were effectively focused into a 1 mm-thick flow cell, where the sample solution was circulated. The circulation rate was meticulously optimized to guarantee the replacement of the excited volume with a fresh solution between successive laser shots, maintaining the integrity of our experimental conditions.

### Data analysis

The data was preprocessed and then subjected to global fitting using KiMoPack.^65^ The approach for global analysis is outlined in the reference.^66^ Evolution-associated difference spectra (EADS) and decay-related difference spectra (DADS) were computed using a Python package, and the analytically derived time constants were applied to both types of spectra. The evolution of intermediate protein states can be readily discerned using sequential models.^54,66^ The standard errors in the time constants were *<* 5% for the transient absorption.

## Acknowledgement

This work is supported by Chinese Technological Innovation 2030- ”Quantum Communication and Quantum Computers” Major Project 3.3. GaoShang Li thanks Pengcheng Mao at the Analysis & Testing Center, Beijing Institute of Technology and Jiayu Wang, Prof. Zhuan Wang, Prof. Yuxiang Weng at Institute of Physics, Chinese Academy of Sciences for their assistance with TA measurements. We acknowledge the Protein Preparation and Characterization Core Facility of Tsinghua University Branch of China National Center for Protein Sciences Beijing for providing the facility support. We thank the Biological and Medical Engineering Core Facilities of Beijing Institute of Technology for supporting experimental equipments. Y.S. acknowledges support from the NSFC (No. 62105030).

## Supporting Information Available

The Supporting Information is available free of charge at https://pubs.acs.org.

## References

(1) Dixon, R. A.; Kobilka, B. K.; Strader, D. J.; Benovic, J. L.; Dohlman, H. G.; Frielle, T.; Bolanowski, M. A.; Bennett, C. D.; Rands, E.; Diehl, R. E.; others Cloning of the gene and cDNA for mammalian *β*-adrenergic receptor and homology with rhodopsin. Nature 1986, 321, 75–79.

(2) Ernst, O. P.; Lodowski, D. T.; Elstner, M.; Hegemann, P.; Brown, L. S.; Kandori, H. Microbial and animal rhodopsins: structures, functions, and molecular mechanisms. Chemical reviews 2014, 114, 126–163.

(3) Oesterhelt, D.; Stoeckenius, W. Rhodopsin-like protein from the purple membrane of Halobacterium halobium. Nature new biology 1971, 233, 149–152.

(4) Mukohata, Y.; Ihara, K.; Uegaki, K.; Mlyashita, Y.; Sugiyama, Y. Australian Halobacteria and their retinal-protein ion pumps. Photochemistry and photobiology 1991, 54, 1039–1045.

(5) Béja, O.; Aravind, L.; Koonin, E. V.; Suzuki, M. T.; Hadd, A.; Nguyen, L. P.; Jovanovich, S. B.; Gates, C. M.; Feldman, R. A.; Spudich, J. L.; others Bacterial rhodopsin: evidence for a new type of phototrophy in the sea. Science 2000, 289, 1902–1906.

(6) Balashov, S. P.; Imasheva, E. S.; Boichenko, V. A.; Antón, J.; Wang, J. M.; Lanyi, J. K. Xanthorhodopsin: a proton pump with a light-harvesting carotenoid antenna. Science 2005, 309, 2061–2064.

(7) Waschuk, S. A.; Bezerra Jr, A. G.; Shi, L.; Brown, L. S. Leptosphaeria rhodopsin: bacteriorhodopsin-like proton pump from a eukaryote. Proceedings of the National Academy of Sciences 2005, 102, 6879–6883.

(8) Miranda, M. R.; Choi, A. R.; Shi, L.; Bezerra, A. G.; Jung, K.-H.; Brown, L. S. The photocycle and proton translocation pathway in a cyanobacterial ion-pumping rhodopsin. Biophysical journal 2009, 96, 1471–1481.

(9) Harris, A.; Ljumovic, M.; Bondar, A.-N.; Shibata, Y.; Ito, S.; Inoue, K.; Kandori, H.; Brown, L. S. A new group of eubacterial light-driven retinal-binding proton pumps with an unusual cytoplasmic proton donor. Biochimica et Biophysica Acta (BBA)- Bioenergetics 2015, 1847, 1518–1529.

(10) Béja, O.; Spudich, E. N.; Spudich, J. L.; Leclerc, M.; DeLong, E. F. Proteorhodopsin phototrophy in the ocean. Nature 2001, 411, 786–789.

(11) Walter, J. M.; Greenfield, D.; Bustamante, C.; Liphardt, J. Light-powering Escherichia coli with proteorhodopsin. Proceedings of the National Academy of Sciences 2007, 104, 2408–2412.

(12) Akram, N.; Palovaara, J.; Forsberg, J.; Lindh, M. V.; Milton, D. L.; Luo, H.; González, J. M.; Pinhassi, J. Regulation of proteorhodopsin gene expression by nutrient limitation in the marine bacterium V ibrio sp. AND 4. Environmental microbiology 2013, 15, 1400–1415.

(13) Kawanabe, A.; Kandori, H. Photoreactions and structural changes of Anabaena sensory rhodopsin. Sensors 2009, 9, 9741–9804.

(14) Jung, K.-H.; Trivedi, V. D.; Spudich, J. L. Demonstration of a sensory rhodopsin in eubacteria. Molecular microbiology 2003, 47, 1513–1522.

(15) Venter, J. C.; Remington, K.; Heidelberg, J. F.; Halpern, A. L.; Rusch, D.; Eisen, J. A.; Wu, D.; Paulsen, I.; Nelson, K. E.; Nelson, W.; others Environmental genome shotgun sequencing of the Sargasso Sea. science 2004, 304, 66–74.

(16) Zabelskii, D.; Alekseev, A.; Kovalev, K.; Rankovic, V.; Balandin, T.; Soloviov, D.; Bratanov, D.; Savelyeva, E.; Podolyak, E.; Volkov, D.; others Viral rhodopsins 1 are an unique family of light-gated cation channels. Nature communications 2020, 11, 1–16.

(17) Bulzu, P.-A.; Andrei, A.-Ş.; Salcher, M. M.; Mehrshad, M.; Inoue, K.; Kandori, H.; Beja, O.; Ghai, R.; Banciu, H. L. Casting light on Asgardarchaeota metabolism in a sunlit microoxic niche. Nature microbiology 2019, 4, 1129–1137.

(18) Inoue, K.; Tsunoda, S. P.; Singh, M.; Tomida, S.; Hososhima, S.; Konno, M.; Nakamura, R.; Watanabe, H.; Bulzu, P.-A.; Banciu, H. L.; others Schizorhodopsins: A family of rhodopsins from Asgard archaea that function as light-driven inward H+ pumps. Science Advances 2020, 6, eaaz2441.

(19) Pushkarev, A.; Inoue, K.; Larom, S.; Flores-Uribe, J.; Singh, M.; Konno, M.; Tomida, S.; Ito, S.; Nakamura, R.; Tsunoda, S. P.; others A distinct abundant group of microbial rhodopsins discovered using functional metagenomics. Nature 2018, 558, 595–599.

(20) Schobert, B.; Lanyi, J. K. Halorhodopsin is a light-driven chloride pump. Journal of Biological Chemistry 1982, 257, 10306–10313.

(21) Matsuno-Yagi, A.; Mukohata, Y. Two possible roles of bacteriorhodopsin; a comparative study of strains of Halobacterium halobium differing in pigmentation. Biochemical and biophysical research communications 1977, 78, 237–243.

(22) Kim, J. E.; Tauber, M. J.; Mathies, R. A. Wavelength dependent cis-trans isomerization in vision. Biochemistry 2001, 40, 13774–13778.

(23) Schoenlein, R.; Peteanu, L.; Mathies, R.; Shank, C. The first step in vision: femtosecond isomerization of rhodopsin. Science 1991, 254, 412–415.

(24) Haran, G.; Morlino, E. A.; Matthes, J.; Callender, R. H.; Hochstrasser, R. M. Femtosecond polarized pump-probe and stimulated emission spectroscopy of the isomerization reaction of rhodopsin. The Journal of Physical Chemistry A 1999, 103, 2202–2207.

(25) Chosrowjan, H.; Mataga, N.; Shibata, Y.; Tachibanaki, S.; Kandori, H.; Shichida, Y.; Okada, T.; Kouyama, T. Rhodopsin emission in real time: a new aspect of the primary event in vision. Journal of the American Chemical Society 1998, 120, 9706–9707.

(26) Zhang, F.; Prigge, M.; Beyrière, F.; Tsunoda, S. P.; Mattis, J.; Yizhar, O.; Hegemann, P.; Deisseroth, K. Red-shifted optogenetic excitation: a tool for fast neural control derived from Volvox carteri. Nature neuroscience 2008, 11, 631–633.

(27) Kandori, H. Biophysics of rhodopsins and optogenetics. Biophysical Reviews 2020, 12, 355–361.

(28) Kandori, H. Retinal proteins: photochemistry and optogenetics. Bulletin of the Chemical Society of Japan 2020, 93, 76–85.

(29) Sivozhelezov, V.; Nicolini, C. Prospects for octopus rhodopsin utilization in optical and quantum computation. Physics of Particles and Nuclei Letters 2007, 4, 189–196.

(30) Kojima, K.; Shibukawa, A.; Sudo, Y. The unlimited potential of microbial rhodopsins as optical tools. Biochemistry 2019, 59, 218–229.

(31) Mahyad, B.; Janfaza, S.; Hosseini, E. S. Bio-nano hybrid materials based on bacteriorhodopsin: potential applications and future strategies. Advances in colloid and interface science 2015, 225, 194–202.

(32) Bakaraju, V.; Prasad, E. S.; Meena, B.; Chaturvedi, H. An Electronic and Optically Controlled Bifunctional Transistor Based on a Bio–Nano Hybrid Complex. ACS omega 2020, 5, 9702–9706.

(33) Boyden, E. S.; Zhang, F.; Bamberg, E.; Nagel, G.; Deisseroth, K. Millisecond-timescale, genetically targeted optical control of neural activity. Nature neuroscience 2005, 8, 1263–1268.

(34) Crist, R. M.; Vasileiou, C.; Rabago-Smith, M.; Geiger, J. H.; Borhan, B. Engineering a rhodopsin protein mimic. Journal of the American Chemical Society 2006, 128, 4522– 4523.

(35) Vaezeslami, S.; Mathes, E.; Vasileiou, C.; Borhan, B.; Geiger, J. H. The structure of apo-wild-type cellular retinoic acid binding protein II at 1.4 Å and its relationship to ligand binding and nuclear translocation. Journal of molecular biology 2006, 363, 687–701.

(36) Vasileiou, C.; Vaezeslami, S.; Crist, R. M.; Rabago-Smith, M.; Geiger, J. H.; Borhan, B. Protein design: reengineering cellular retinoic acid binding protein II into a rhodopsin protein mimic. Journal of the American Chemical Society 2007, 129, 6140–6148.

(37) Wang, W.; Nossoni, Z.; Berbasova, T.; Watson, C. T.; Yapici, I.; Lee, K. S. S.; Vasileiou, C.; Geiger, J. H.; Borhan, B. Tuning the electronic absorption of protein-embedded all-trans-retinal. Science 2012, 338, 1340–1343.

(38) Lee, K. S. S.; Berbasova, T.; Vasileiou, C.; Jia, X.; Wang, W.; Choi, Y.; Nossoni, F.; Geiger, J. H.; Borhan, B. Probing Wavelength Regulation with an Engineered Rhodopsin Mimic and a C15-Retinal Analogue. ChemPlusChem 2012, 77, 273–276.

(39) Berbasova, T.; Nosrati, M.; Vasileiou, C.; Wang, W.; Lee, K. S. S.; Yapici, I.; Geiger, J. H.; Borhan, B. Rational design of a colorimetric pH sensor from a soluble retinoic acid chaperone. Journal of the American Chemical Society 2013, 135, 16111–16119.

(40) Wang, W.; Geiger, J. H.; Borhan, B. The photochemical determinants of color vision: revealing how opsins tune their chromophore’s absorption wavelength. Bioessays 2014, 36, 65–74.

(41) Yapici, I.; Lee, K. S. S.; Berbasova, T.; Nosrati, M.; Jia, X.; Vasileiou, C.; Wang, W.; Santos, E. M.; Geiger, J. H.; Borhan, B. “Turn-on” protein fluorescence: In situ formation of cyanine dyes. Journal of the American Chemical Society 2015, 137, 1073–1080.

(42) Nosrati, M.; Berbasova, T.; Vasileiou, C.; Borhan, B.; Geiger, J. H. A Photoisomerizing Rhodopsin Mimic Observed at Atomic Resolution. Journal of the American Chemical Society 2016, 138, 8802–8808.

(43) Huntress, M. M.; Gozem, S.; Malley, K. R.; Jailaubekov, A. E.; Vasileiou, C.; Vengris, M.; Geiger, J. H.; Borhan, B.; Schapiro, I.; Larsen, D. S.; others Toward an understanding of the retinal chromophore in rhodopsin mimics. The Journal of Physical Chemistry B 2013, 117, 10053–10070.

(44) Manathunga, M.; Jenkins, A. J.; Orozco-Gonzalez, Y.; Ghanbarpour, A.; Borhan, B.; Geiger, J. H.; Larsen, D. S.; Olivucci, M. Computational and spectroscopic characterization of the photocycle of an artificial rhodopsin. The journal of physical chemistry letters 2020, 11, 4245–4252.

(45) Demoulin, B.; Maiuri, M.; Berbasova, T.; Geiger, J. H.; Borhan, B.; Garavelli, M.; Cerullo, G.; Rivalta, I. Control of protonated Schiff base excited state decay within visual protein mimics: a unified model for retinal chromophores. Chemistry–A European Journal 2021, 27, 16389–16400.

(46) Li, G.; Hu, Y.; Pei, S.; Meng, J.; Wang, J.; Wang, J.; Yue, S.; Wang, Z.; Wang, S.; Liu, X.; others Excited-state dynamics of all-trans protonated retinal Schiff base in CRABPII-based rhodopsin mimics. Biophysical Journal 2022, 121, 4109–4118.

(47) Ghanbarpour, A.; Nairat, M.; Nosrati, M.; Santos, E. M.; Vasileiou, C.; Dantus, M.; Borhan, B.; Geiger, J. H. Mimicking microbial rhodopsin isomerization in a single crystal. Journal of the American Chemical Society 2018, 141, 1735–1741.

(48) Wand, A.; Friedman, N.; Sheves, M.; Ruhman, S. Ultrafast photochemistry of light-adapted and dark-adapted bacteriorhodopsin: effects of the initial retinal configuration. The Journal of Physical Chemistry B 2012, 116, 10444–10452.

(49) Wand, A.; Rozin, R.; Eliash, T.; Jung, K.-H.; Sheves, M.; Ruhman, S. Asymmetric toggling of a natural photoswitch: Ultrafast spectroscopy of Anabaena sensory rhodopsin. Journal of the American Chemical Society 2011, 133, 20922–20932.

(50) Roy, P. P.; Kato, Y.; Abe-Yoshizumi, R.; Pieri, E.; Ferré, N.; Kandori, H.; Buckup, T. Mapping the ultrafast vibrational dynamics of all-trans and 13-cis retinal isomerization in Anabaena Sensory Rhodopsin. Physical Chemistry Chemical Physics 2018, 20, 30159–30173.

(51) Malakar, P.; Das, I.; Bhattacharya, S.; Harris, A.; Sheves, M.; Brown, L. S.; Ruhman, S. Bidirectional Photochemistry of Antarctic Microbial Rhodopsin: Emerging Trend of Ballistic Photoisomerization from the 13-cis Resting State. The journal of physical chemistry letters 2022, 13, 8134–8140.

(52) Roy, P. P.; Youshizumi, R.; Kandori, H.; Buckup, T. Mapping the ultrafast vibrational dynamics of all-trans and 13-Cis retinal isomerization in Anabaena Sensory Rhodopsin. EPJ Web of Conferences. 2019; p 10001.

(53) Ruckebusch, C.; Sliwa, M.; Pernot, P. d.; De Juan, A.; Tauler, R. Comprehensive data analysis of femtosecond transient absorption spectra: A review. Journal of Photochemistry and Photobiology C: Photochemistry Reviews 2012, 13, 1–27.

(54) Snellenburg, J. J.; Laptenok, S.; Seger, R.; Mullen, K. M.; van Stokkum, I. H. Glotaran: A Java-based graphical user interface for the R package TIMP. Journal of Statistical Software 2012, 49, 1–22.

(55) Johnson, P. J.; Halpin, A.; Morizumi, T.; Brown, L. S.; Prokhorenko, V. I.; Ernst, O. P.; Miller, R. D. The photocycle and ultrafast vibrational dynamics of bacteriorhodopsin in lipid nanodiscs. Physical Chemistry Chemical Physics 2014, 16, 21310–21320.

(56) Lenz, M. O.; Huber, R.; Schmidt, B.; Gilch, P.; Kalmbach, R.; Engelhard, M.; Wachtveitl, J. First steps of retinal photoisomerization in proteorhodopsin. Biophysical journal 2006, 91, 255–262.

(57) Tahara, S.; Takeuchi, S.; Abe-Yoshizumi, R.; Inoue, K.; Ohtani, H.; Kandori, H.; Tahara, T. Ultrafast photoreaction dynamics of a light-driven sodium-ion-pumping retinal protein from Krokinobacter eikastus revealed by femtosecond time-resolved absorption spectroscopy. The Journal of Physical Chemistry Letters 2015, 6, 4481–4486.

(58) Schapiro, I.; Ruhman, S. Ultrafast photochemistry of anabaena sensory rhodopsin: Experiment and theory. Biochimica et Biophysica Acta (BBA)-Bioenergetics 2014, 1837, 589–597.

(59) Cheminal, A.; Léonard, J.; Kim, S.-Y.; Jung, K.-H.; Kandori, H.; Haacke, S. 100 fs photo-isomerization with vibrational coherences but low quantum yield in Anabaena Sensory Rhodopsin. Physical Chemistry Chemical Physics 2015, 17, 25429–25439.

(60) Liebel, M.; Schnedermann, C.; Bassolino, G.; Taylor, G.; Watts, A.; Kukura, P. Direct observation of the coherent nuclear response after the absorption of a photon. Physical review letters 2014, 112, 238301.

(61) Stensitzki, T.; Muders, V.; Schlesinger, R.; Heberle, J.; Heyne, K. The primary photoreaction of channelrhodopsin-1: wavelength dependent photoreactions induced by ground-state heterogeneity. Frontiers in Molecular Biosciences 2015, 2, 41.

(62) Tahara, S.; Singh, M.; Kuramochi, H.; Shihoya, W.; Inoue, K.; Nureki, O.; Béjà, O.; Mizutani, Y.; Kandori, H.; Tahara, T. Ultrafast dynamics of heliorhodopsins. The Journal of Physical Chemistry B 2019, 123, 2507–2512.

(63) Inoue, K.; Tahara, S.; Kato, Y.; Takeuchi, S.; Tahara, T.; Kandori, H. Spectroscopic study of proton-transfer mechanism of inward proton-pump rhodopsin, Parvularcula oceani xenorhodopsin. The Journal of Physical Chemistry B 2018, 122, 6453–6461.

(64) Lu, H.; Chen, K.; Bobba, R. S.; Shi, J.; Li, M.; Wang, Y.; Xue, J.; Xue, P.; Zheng, X.; Thorn, K. E.; others Simultaneously Enhancing Exciton/Charge Transport in Organic Solar Cells by an Organoboron Additive. Advanced Materials 2022, 34, 2205926.

(65) Müller, C.; Pascher, T.; Eriksson, A.; Chabera, P.; Uhlig, J. KiMoPack: A python Package for Kinetic Modeling of the Chemical Mechanism. The Journal of Physical Chemistry A 2022, 126, 4087–4099.

(66) van Stokkum, I. H.; Larsen, D. S.; Van Grondelle, R. Global and target analysis of time-resolved spectra. Biochimica et Biophysica Acta (BBA)-Bioenergetics 2004, 1657, 82–104.

